# Functional Connectivity Alterations in Spinocerebellar Ataxia Type 10: Insights from Gray Matter Atrophy

**DOI:** 10.1101/2025.03.19.644149

**Authors:** Gustavo Padron-Rivera, Gabriel Ramirez-Garcia, Amanda Chirino-Perez, Angel Omar Romero-Molina, Adriana Ochoa-Morales, María Guadalupe Garcia-Gomar, Miguel Angel Ramirez-Garcia, Omar Rodriguez-Mendoza, Diana Laura Torres, Birgitt Schüle, Erick Humberto Pasaye-Alcaraz, Carlos Roberto Hernandez-Castillo, Juan Fernandez-Ruiz

**Affiliations:** Laboratorio de Neuropsicologia, Departamento de Fisiologia, Facultad de Medicina, Universidad Nacional Autonoma de Mexico, Ciudad de Mexico, Mexico; Departamento de Neurogenetica, Instituto Nacional de Neurologia y Neurocirugia Manuel Velasco Suarez, Ciudad de Mexico, Mexico; Escuela Nacional de Estudios Superiores (ENES), Universidad Nacional Autónoma de México, Juriquilla Queretaro, Mexico; Department Pathology, Stanford University School of Medicine, Stanford, California, USA; Unidad de Resonancia Magnetica, Instituto de Neurobiología, Universidad Nacional Autónoma de México, Juriquilla Queretaro, Mexico; Faculty of Computer Science, Dalhousie University, Halifax, Canada; Instituto de Neuroetologia Universidad Veracruzana, Xalapa Veracruz, Mexico

**Keywords:** SCA10, functional connectivity, grey matter atrophy, cerebellum, motor cortex

## Abstract

Spinocerebellar ataxia type 10 (SCA10) is a rare, inherited neurological disorder caused by an expansion of the non-coding ATTCT pentanucleotide repeat in the ATAXIN 10 gene. It is characterized by cerebellar ataxia and epilepsy. Previous research has demonstrated extensive white and gray matter degeneration, particularly in the cerebellum. However, the impact of the SCA10 mutation on functional connectivity (FC) remains unexplored. This study aimed to characterize intrinsic FC changes in SCA10 patients and their relationship to clinical manifestations. Structural and resting-state MRIs were obtained from 26 SCA10 patients and 26 control subjects. Voxel-based morphometry (VBM) and seed-ROI and Independent Components Analysis (ICA) were performed to identify cerebral atrophy and FC changes respectively. Additionally, correlation analyses were conducted between FC changes and scores from the Scale for the Assessment and Rating of Ataxia (SARA) and the Montreal Cognitive Assessment (MoCA). In SCA10 patients, VBM analysis revealed extensive gray matter loss in motor cortices and the cerebellum. FC analysis identified significant FC changes originating from seed-ROIs in the right cerebellar VI and left precentral gyrus. Furthermore, group comparison using ICA components showed that SCA10 patients exhibited higher FC in the sensorimotor and cerebellar functional networks. Moreover, the average BOLD signal within the cerebellar network negatively correlated with MoCA scores. In summary, SCA10 patients exhibited enhanced FC in brain regions that displayed gray matter atrophy, underscoring the impact of SCA10 degeneration on resting state networks and induction of potential maladaptive FC compensatory mechanisms.

## Introduction

Spinocerebellar ataxia type 10 (SCA10) is a unique autosomal dominant disorder characterized by the co-occurrence of cerebellar ataxia and epilepsy. This complex clinical profile arises from the expansion of a pentanucleotide repeat sequence (ATTCT) within the ATXN10 gene [1]. In addition to its core features, SCA10 is associated with a range of additional symptoms, including polyneuropathy, pyramidal signs, dysarthria, as well as cognitive and neuropsychiatric impairments [2] [3]. Understanding the full clinical spectrum of this SCA subtype requires a comprehensive characterization of the structural and functional brain changes in affected individuals. However, due to the rarity of SCA10, limited reports have investigated these changes compared to other types of SCAs [4]. Structural studies in SCA10 patients have revealed extensive cerebellar atrophy in both white and gray matter, along with moderate cortical, pallidal, brainstem, thalamic, and putaminal degeneration [5] [6]. While initial studies have provided anatomical insights into brain atrophy in SCA10, there is no information regarding functional connectivity (FC) alterations and their possible relationship to clinical metrics.

Therefore, this study aimed to address the following objectives: 1) investigate the FC changes of resting-state networks (RSNs) associated with grey matter atrophy nodes. 2) explore potential alterations in the FC of distinctive RSNs, and 3) determine correlations between the BOLD signal of both atrophy-based and distinctive RSNs and cognitive and ataxia assessments. By addressing these objectives, we seek to provide a deeper understanding of the neural mechanisms underlying SCA10 and their relationship to clinical manifestations, ultimately contributing to improved diagnostic and therapeutic strategies for this challenging condition.

## Methods

### Participants

Twenty-six patients of SCA10 were included in this study of whom three were asymptomatic individuals with molecular diagnosis confirming their condition. Twenty-six right-handed subjects were selected as controls, matched to the SCA10 group in terms of age, gender, and level of education (see Table 1). All procedures were approved by the ethics committee of the Universidad Nacional Autónoma de México (UNAM) following the principles outlined in the Helsinki Declaration. Before participation, written consent was obtained from each participant.

**Table 1.** Demographic and clinical characteristics of the participants.

|  | All |  |  | fMRI |  |  |
| --- | --- | --- | --- | --- | --- | --- |
|  | Control | SCA10 | p-value/t-value | Control | SCA10 | p-value/t-value |
| n | 26 | 26 | – | 20 | 19 | – |
| Male/Female | 11 / 15 | 11 / 15 | – | 8 / 13 | 7 / 12 | – |
| Age | 50.65 ± 9.28 | 50.38 ± 9.91 | 0.922/<br>t=0.098 | 50.56 ± 8.13 | 50.10 ± 9.8 | 0.876/<br>t=0.157 |
| Year of education | 10.15 ± 6 | 9.73 ± 2.50 | 0.742/<br>t=0.331 | 9.05 ± 6.13 | 10.11 ± 2.58 | 0.492/<br>t=0.693 |
| Age of onset | – | 33.82 ± 8.66 | – | – | 34.62 ± 8.56 | – |
| Disease duration | – | 16.17 ± 9.57 | – | – | 14.87 ± 10.34 | – |
| SARA | – | 15.7 ± 7.97 | – | – | 14.4 ± 8.13 | – |
| MoCA | 27.0 ± 2.42 | 21.9 ± 4.15 | 0.001***/<br>t=4.275 | 26.64 ± 2.58 | 21.3 ± 4.27 | 0.0009***/<br>t=3.702 |

\*Values represent mean ± SD

### Clinical assessment

The Scale for the Assessment and Rating of Ataxia (SARA) was utilized to evaluate the fundamental symptoms of ataxia in the participants. This concise clinical score consists of 8 items assessing various aspects including gait, stance, sitting, speech disturbance, finger-chase, nose-finger test, fast alternating hand movements, and heel-shin slide. Scores on the SARA range up to a maximum of 40 points, with higher scores indicating more severe ataxia [7]. Additionally, the Montreal Cognitive Assessment (MoCA) Spanish version 1 was employed as a cognitive screening measure. This assessment evaluates executive function, verbal memory, visuospatial ability, attention, working memory, language, abstract reasoning, and orientation to time and place. The MoCA total score, with a maximum of 30 points, provides an overall assessment of cognitive performance. A score equal to or below 25 points is indicative of mild cognitive impairment [8] [9].

### Image Acquisition

The imaging procedures were conducted at the Instituto de Neurobiologia of UNAM in Juriquilla, Queretaro, Mexico, utilizing a 3T General Electric MR750 Discovery scanner equipped with a 32-channel head coil. A high-resolution T1-3D volume was acquired with a TR/TE of 3.18/8.16 ms, flip angle of 9°, and FOV and matrix of 256 × 256, resulting in an isometric resolution of 1 × 1 × 1 mm³. Resting-state functional images were obtained using an Echo Planar Imaging single-shot sequence with a TR/TE of 2000/35 ms, FOV of 212 × 212 mm², flip angle of 80°, and 150 whole-brain volumes comprising 91 slices each. The final resolution was 3 × 3 × 4 mm, with no gaps between slices.

### Voxel-Based Morphometry Analysis to Determine Structural Atrophy Nodes

The analysis utilized whole-brain Voxel-based Morphometry (VBM) to identify changes in gray matter (GM) volume between groups. FSL-VBM (https://fsl.fmrib.ox.ac.uk/fsl/docs/#/structural/fslvbm) [13] was employed, which includes skull stripping, segmentation into gray matter (GM), white matter, and cerebrospinal fluid. Subsequently, GM images were normalized to MNI152 standard space, modulated, and smoothed. A two-sample t-test, utilizing the randomize tool with 10,000 permutations [14], was conducted to compare SCA10 patients and Control subjects. Age served as a nuisance regressor, and maps were thresholded at p < 0.05 and corrected by Threshold-Free Cluster Enhancement (TFCE) for family-wise errors. Anatomical descriptions of GM changes were referenced using the Harvard-Oxford Cortical Structural Atlas, the Harvard-Oxford Subcortical Structural Atlas, and the Cerebellar Atlas in MNI152.

Following this analysis, a set of regions of interest (ROIs) were defined based on the highest local maxima peak values within the significant clusters obtained. Each ROI was centered at the local maxima voxel within the clusters and consisted of a 12-mm sphere size.

### Resting-state fMRI analysis

The preprocessing of functional images involved several steps: motion and slice timing correction, removal of non-brain tissue, spatial smoothing using a 5-mm full-width-at-half-maximum Gaussian kernel, and high-pass temporal filtering equivalent to 100s (0.01 Hz), in order to remove slow drifts. Also, the resting state signal is low frequency, mostly between 0.01-0.1 Hz, consequently, we want to remove frequencies only below 0.01 Hz (corresponding to a period of 100 s). Additionally, we relied on previous research for preprocessing rsfMRI data to identify intrinsic brain activity [10] [11]. Following preprocessing, the resting-state volumes were registered first to the subject’s high-resolution T1-weighted scan and then to the standard MNI152 space using nonlinear registration. Subsequently, ICA-AROMA was applied to effectively eliminate motion-related spurious noise from the resting-state fMRI datasets [12]. After preprocessing, seven participants were discarded due to motion artifacts across volumes that affected subsequent analysis. The final sample size for the rsfMRI analysis included 19 SCA10 patients, who were compared to 20 Control subjects.

### Independent Components Analysis and Dual regression

Independent Components Analysis (ICA) was conducted using MELODIC (Multivariate Exploratory Linear Decomposition into Independent Components) within FSL. Preprocessed volumes were initially analyzed on a single-subject basis. Twenty-five components were extracted per subject, and manual classification of these components was conducted. Components corresponding to resting-state networks (RSNs) were selected for further analysis, while components identified as structured noise were removed through regression-based denoising. The selected RSNs from each subject were then analyzed and grouped using a temporal-concatenation approach, which was based on the frequency spectra of the components’ time courses. The compiled dataset was ultimately decomposed into 25 independent components [13]. To identify RSNs in this study, single ICs were projected on the resting state functional network atlas (available at http://www.fmrib.ox.ac.uk/analysis/royalsoc8/) and visual inspections of previously reported RSNs were performed [14] [15] [16] [17] [18] [19] [20].

To compare RSNs between groups, a dual-regression analysis was employed [21]. This method utilizes the ICs maps as network templates to identify corresponding functional connectivity maps for each subject [22], integrating temporal information in the resting state fMRI data across multiple distributed networks identified in the initial group ICA [21]. A two-sample t-test with 10,000 permutations was then conducted and corrected by TFCE to determine significant differences between groups. Age was included as a nuisance regressor in the analysis

### Seed-Based Functional Connectivity Analysis

Four seeds-ROI were defined from the VBM analysis. A voxel-wise seed-based analysis was conducted by computing Pearson’s correlation between the 4D denoised data and each ROI. The correlation values were then normalized into Fisher’s Z-score. Functional maps obtained from each ROI were compared between SCA10 patients and Control subjects using a two-sample t-test with 10,000 permutations. Age was included as a nuisance regressor in the analysis. The maps were thresholded at p < 0.05, with correction using TFCE. This analysis was performed using MATLAB R2015b (The MathWorks Inc., Natick, MA).

### Statistical Analysis

Spearman’s partial correlations were calculated between the average BOLD signal of all clusters within each RSN that showed significant differences between groups and the MoCA and SARA scores, controlled by age. This correlation analysis included RSNs obtained from both seed-ROI and ICA approaches. Subsequently, an FDR correction for multiple comparisons was applied, setting a significant threshold of p < 0.05. The anatomical descriptions of VBM, ICA and seed-based FC were based on the Harvard-Oxford cortical and subcortical structural atlases and the cerebellar atlas [23]. All statistical analyses were performed with R version 4.4.0.

## Results

Demographic data did not show a significant difference between SCA10 and Control subjects for variables such as sex, age, and years of education. In addition, there were no significant differences in the same variables for the subsample of participants included in the fMRI analysis (Table 1). However, significant differences were found for MoCA scores considering the whole and partial sample, as reported previously [24].

### Voxel-based morphometry analysis

VBM analysis was performed to determine the GM degeneration in the brain and cerebellum in SCA10 patients. SCA10 patients showed a GM decrease in the cortical mantle encompassing the bilateral precentral gyrus, bilateral postcentral gyrus, bilateral supplementary motor area, temporal pole, and partially the superior frontal gyrus, occipital fusiform gyrus, and the temporal occipital fusiform cortex as well as over the whole cerebellum, especially regions such as bilateral Crus I, Crus II, V, VI, VIIIa, VIIb, VIIIb, and right IX (Fig. 1).

**Fig. 1.**
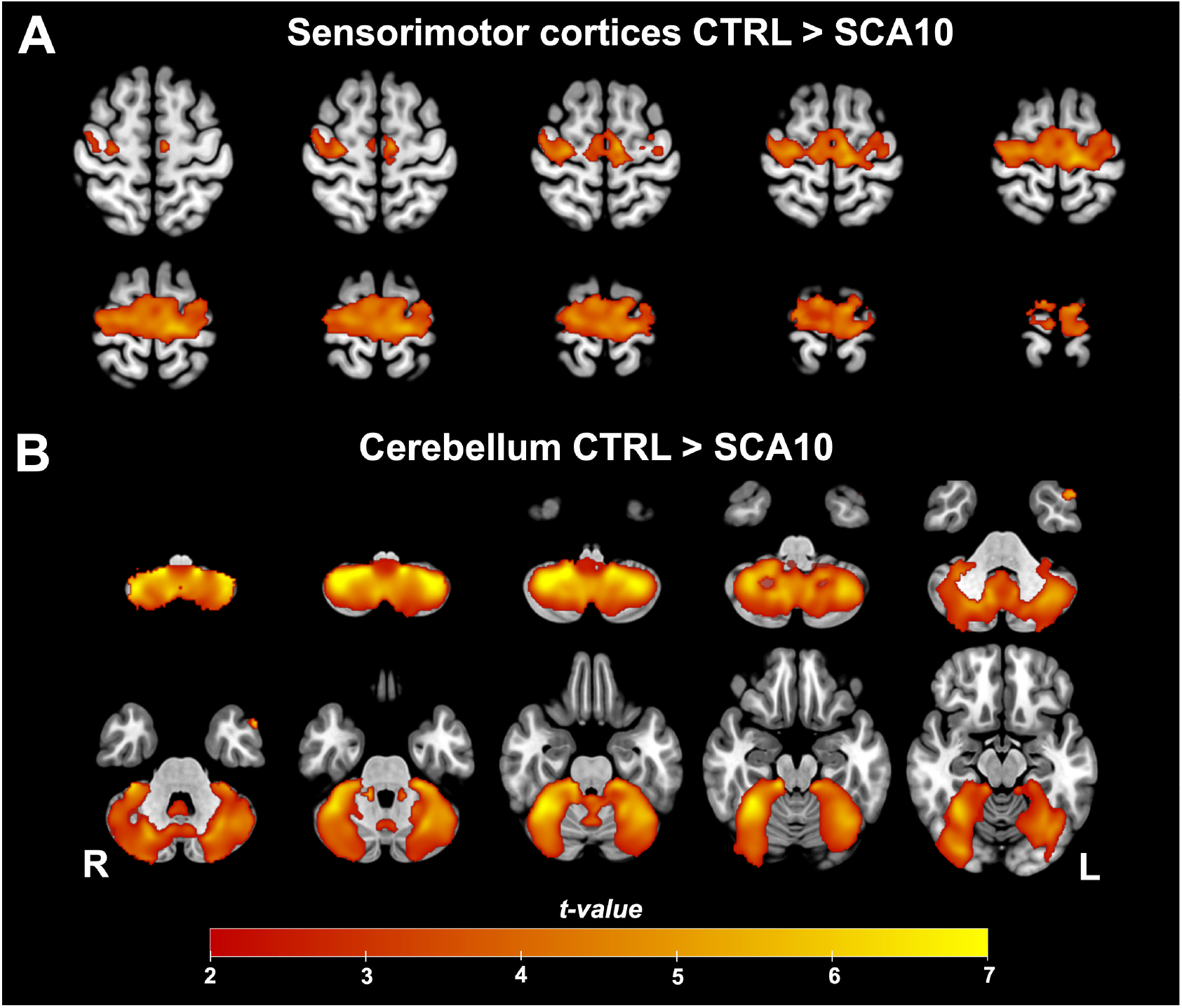
Grey matter Volume reduction in SCA10 patients. Two main clusters were found in the sensorimotor (A) cortices and the cerebellum (B). Warm colors show the areas where SCA10 patients’ GM decreased significantly compared to Controls (p < 0.05)

### ROIs-based Analysis

VBM found two large clusters displaying GM decreases (Table 2). Four spheric ROIs were extracted from these two clusters based on the highest local maxima within them. Regarding the ROIs from the cerebellum, SCA10 patients showed increased FC between right cerebellar VI with precuneus cortex and posterior division of the cingulate gyrus (Fig. 2A). For the seeds located in the right Cerebellum VIIIb and left Cerebellum V no significant differences were found. For the seed located in the left precentral gyrus, significant FC changes were found with the parietal operculum cortex, right precentral and postcentral gyrus, supramarginal gyrus (anterior and posterior division), angular gyrus, precuneus cortex, planum temporalis, parietal operculum, and caudate (Fig. 2B).

**Table 2.** Local maxima coordinate of the seed-ROIs.

| **Anatomical Region** | **MNI coordinates (voxels)** |  |  |
| --- | --- | --- | --- |
|  | **X** | **Y** | **Z** |
| **Cerebellum** |  |  |  |
| Right Cerebellum VIIIb | 36 | 40 | 11 |
| Right Cerebellum VI | 27 | 37 | 19 |
| Left Cerebellum V | 57 | 46 | 22 |
| **Brain** |  |  |  |
| Left Precentral gyrus | 51 | 50 | 70 |

**Fig. 2.**
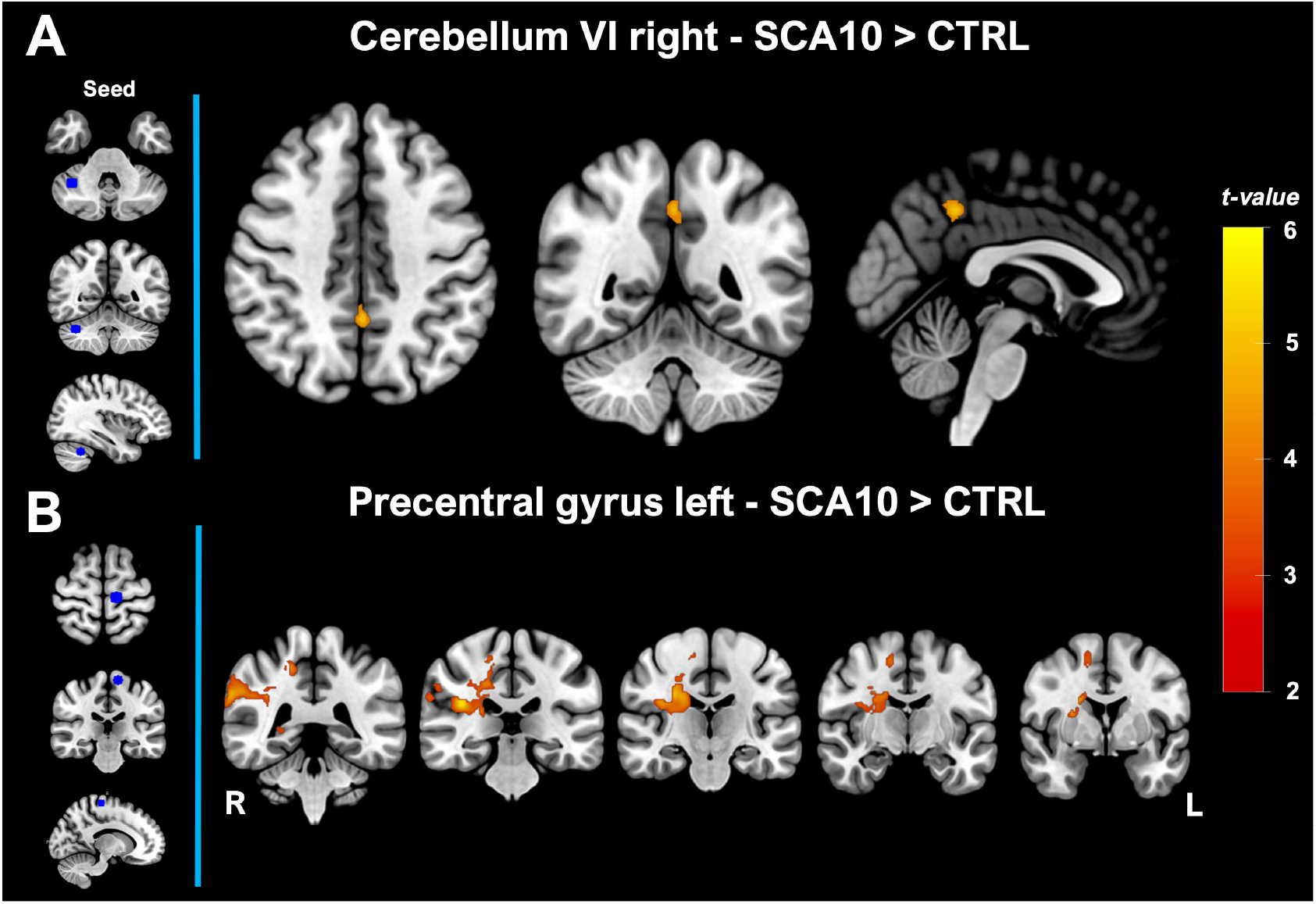
Functional connectivity of seed atrophy based in SCA10 patients. A) shows the FC change for the seed in the cerebellum VI right. B) shows the FC change for the seed in the precentral gyrus left. The left vertical panel showed the location of the spheric seed in blue. The red color bar represents the significant increment of FC in patients with SCA10 (p < 0.05)

### Independent Components Analysis

Besides the seed-ROI-based FC analyses, we also explored possible FC changes using ICA. Sixteen resting-state functional networks were identified encompassing default mode network, executive control network, medial visual network, lateral visual network, occipital visual network, somatosensory network, superior frontal network, somatosensory motor network, auditory network, executive frontal network, frontoparietal network, salience network, cerebellar network, basal ganglia network and prefrontal network (See supplementary data, Fig. 1). Group comparison between these resting state networks showed that SCA10 patients had higher FC in the sensorimotor network and cerebellar network. Especially, the FC increased in the cerebellar network and was grouped in two clusters located in the bilateral Crus I, Crus II, I-IV, V, VI, VIIb, Vermis VI, and Vermis Crus II. In addition, the sensorimotor network showed an FC increase in three clusters located in the posterior division of the cingulate gyrus, precuneus cortex, precentral gyrus, postcentral gyrus, anterior and posterior division of the supramarginal gyrus, angular gyrus, and superior division of the lateral occipital cortex (Fig. 3).

**Fig. 3.**
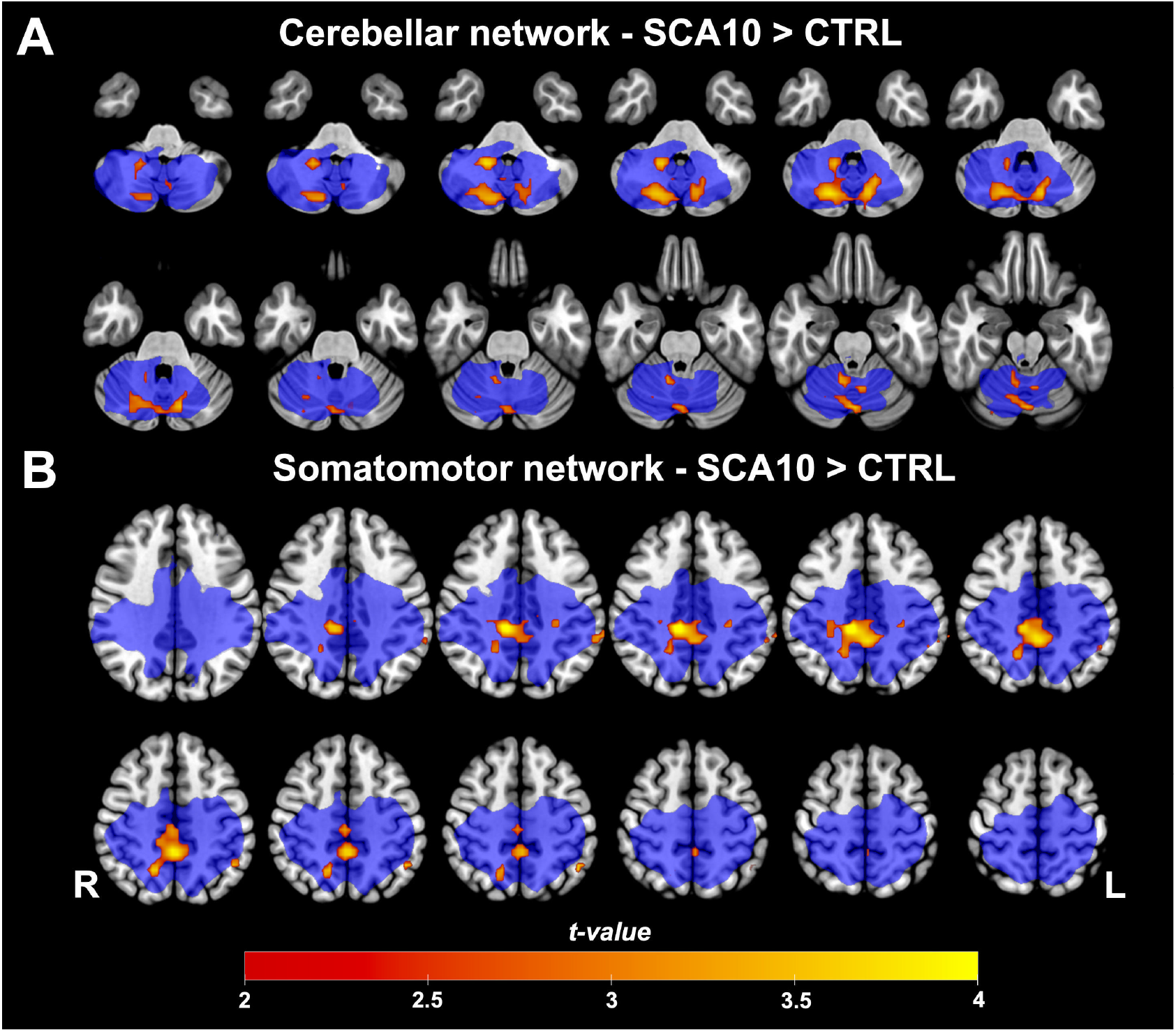
Group differences in cerebellar and somatomotor functional networks. (A) Cerebellar Network and (B) Somatomotor Network compared to Controls. Warm colors represent where SCA10 patients showed significantly increased functional connectivity within the network (p < 0.05. The blue color represents the size and location of the original functional network

### Correlation analysis

The partial correlation analysis of the BOLD signal of the significant RSNs, obtained from both the seed-ROI and ICA analysis, showed that only the Cerebellar network had a significant negative relationship with MoCA scores; r = −0.5293491, p = 0.0237, p-FDR = 0.047 (Fig. 4). There were no significant correlations between any BOLD signal of RSNs and SARA scores.

**Fig. 4.**
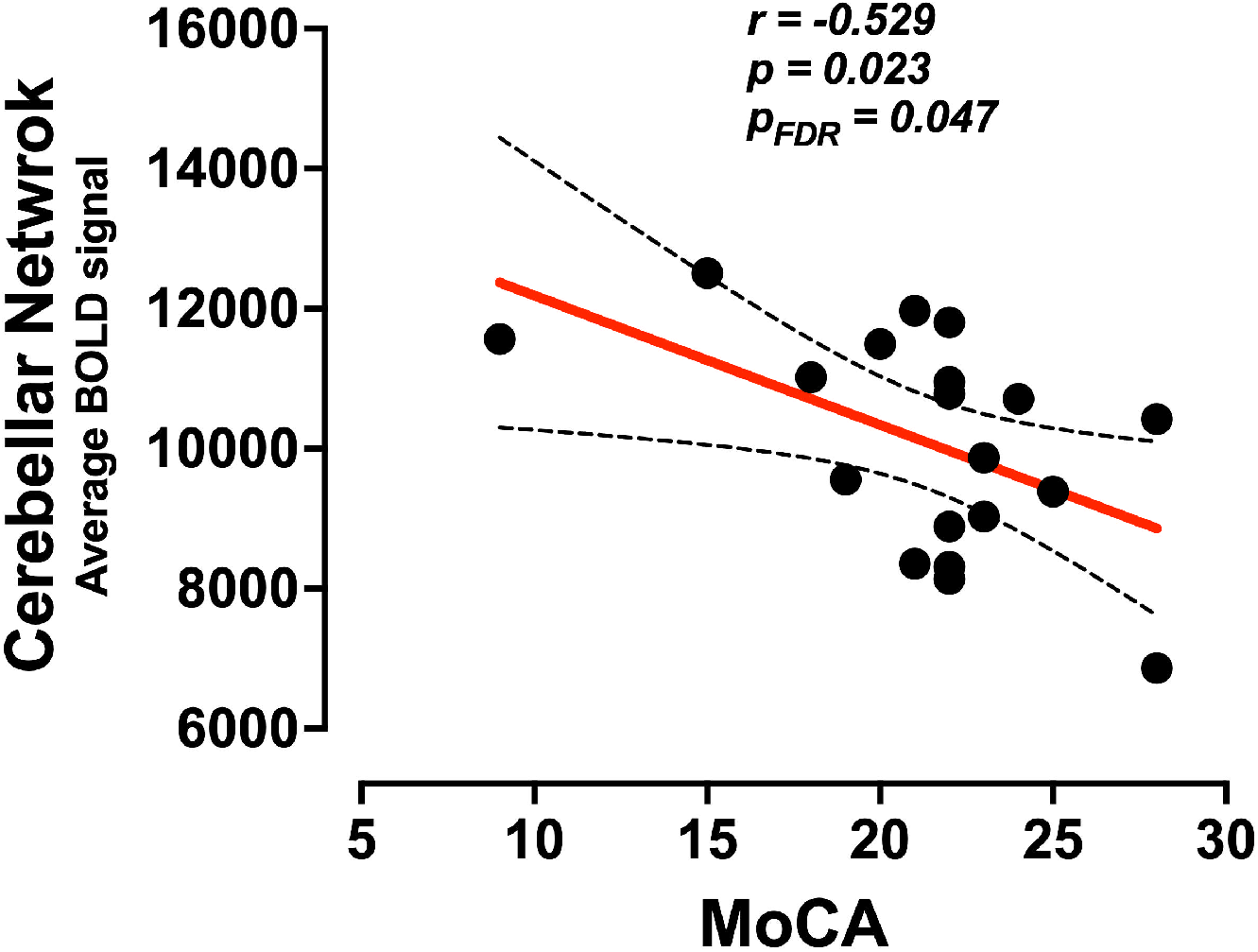
Partial correlation between the Cerebellar network and MoCA scores. Spearman partial correlation between the BOLD signal of the cerebellar network and the MoCA scores (p< 0.05)

## Discussion

SCA10 is a neurodegenerative disease characterized by structural and functional brain damage. Using multimodal MRI, we explored cerebral and cerebellar gray matter atrophy and functional connectivity changes in RSNs associated with GM atrophy. We identified bilateral GM reduction in the cerebellum and motor cortices. The most significant peaks within these atrophy clusters were located in the right cerebellum VI and the left precentral gyrus. Additionally, these ROIs showed higher FC changes in SCA10 patients. Moreover, the canonical cerebellar and somatomotor functional networks exhibited increased FC connectivity in SCA10 patients as well. Lastly, the BOLD signal of the cerebellar network correlated negatively with the cognitive MoCA performance in these patients. These findings provide a comprehensive characterization of alterations in spontaneous resting state networks linked to gray matter atrophy patterns in SCA10 patients. This study represents a founding functional description of SCA10, offering insights into the neurobiological underpinnings of the disease.

SCA10 is a genetic disease primarily affecting the cerebellum. In our study, we found that the cerebellum and the motor and sensorimotor cortices were the two main affected regions. Previous studies in the Brazilian SCA10 population showed extensive cerebellar atrophy along with pallidal reduction. Cortical thickness reduction was observed in the left frontal (frontal pole, orbital middle frontal, and rostral anterior cingulate), temporal (bilateral parahippocampal gyrus), and left occipital (lingual gyrus) lobes. In previous research, with a similar sample size in a Mexican population, the atrophy pattern included the cerebellum, brain stem, thalamus, putamen, and cingulate and precentral gyrus [25] [6]. Despite slight differences in atrophy patterns, GM reduction cores remain in the same anatomical areas: the cerebellum and sensorimotor cortices.

In SCA10 patients, we identified an atrophy core located in the right cerebellum VI, which exhibited higher functional connectivity with the precuneus cortex and the posterior division of the cingulate gyrus. The precuneus is critically involved in higher-order cognitive processes such as visuospatial imagery, episodic memory retrieval, and self-processing operations [26], while the posterior division of the cingulate gyrus is associated with emotional regulation, memory processing, and internally directed thought. This increased connectivity may suggest a compensatory mechanism within the brain’s networks due to cerebellar degeneration. The precuneus and posterior division of the cingulate gyrus are key nodes within the DMN, which is active during rest and involved in internally directed cognitive activities. In our study, we did not find any significant changes in the DMN network as has been reported in SCA3 [27], which represents that the functional integrity of this network remains unaffected at this point in these patients. Additionally, the observed higher functional connectivity between the cerebellum and cingulate gyrus may reflect the cerebellum’s broader role in integrating cognitive and emotional information [28].

A comparison of canonical spontaneous resting state networks showed higher FC in the sensorimotor network. This greater FC may imply an enhanced interaction between motor and sensory processing areas, possibly as a response to cerebellar dysfunction. On the other hand, the higher FC displayed in the cerebellar networks may suggest extensive involvement of the cerebellum in compensatory mechanisms or network reorganization in response to disease-related atrophy as has been shown in SCA2 [29].

These findings indicate that SCA10 not only affects motor coordination but also impacts broader networks associated with sensory processing and cognitive functions. The increased connectivity within these networks may reflect a compensatory process aimed at maintaining functional integrity despite ongoing neurodegeneration. These findings provide valuable insights into the neural mechanisms underlying motor impairments associated with SCA10, suggesting a disrupted interplay between the cerebellum and motor-related brain regions [24].

Finally, we found a correlation between the FC of the cerebellum network and the total MoCA scores per patient. Although the cerebellum has traditionally been described as a regulator of motor actions, recent research suggests that it also controls cognitive and affective behaviors. Specifically, our study revealed a negative correlation between MoCA scores and the average BOLD signal within the cerebellar network. Besides, this finding underscores the cerebellum’s significant role in a variety of behavioral and cognitive domains. In this regard, we suggest that the increased FC observed in resting-state networks may represent a maladaptive compensatory mechanism triggered by atrophy, as evidenced by the patients’ impaired cognitive and motor performance. A previous study described significant memory and executive dysfunction in SCA10 patients, which could be correlated with deterioration in the posterior lobe of the cerebellum, as well as the prefrontal, cingulate, and middle temporal cortices [24]. However, further analyses are needed to fully understand these changes and their potential consequences on the motor and cognitive performance of the patients. To gain a more comprehensive understanding of cognitive alterations in SCA10 patients, incorporating more specific assessments is crucial. The Schmahmann Scale [30], which is specifically designed for the assessment of cerebellar cognitive affective syndrome (CCAS) could provide deeper insights into the specific cognitive deficits associated with cerebellar dysfunction in SCA10, to evaluate executive, linguistic, visuospatial and affective impairments.

## Conclusion

In conclusion, our study offers valuable insights into the neurobiological underpinnings of SCA10 by revealing specific network-level functional connectivity changes linked to cerebellar atrophy. These findings highlight the significant impact of SCA10 degeneration on resting-state networks and suggest the induction of potential maladaptive functional connectivity compensatory mechanisms.

## Acknowledgment

We sincerely thank all the participants, especially the patients who took part in this study, as well as their families for their invaluable support. We also extend our gratitude to the MRI staff (INB, UNAM) for their expertise and dedication in facilitating the scanning process, which was crucial for the success of this research.

## Ethical Approval

The study was performed in accordance with the principles stated in the Declaration of Helsinki and with the ethics committee of the Universidad Nacional Autonoma de Mexico (UNAM).

## Consent to Participate

Informed consent for data publication was obtained from all participants involved in this study.

## Data Availability

The data and material supporting the findings of this study are available upon reasonable request from the corresponding author.

## Author Contributions Statement

Conceptualization: G.PR., G.RG., J.FR.; Methodology: G.PR, G.RG., C.R.HC., J.FR.; Formal analysis and investigation: G.PR., G.RG., J.FR., E.H.PA.; Writing - original draft preparation: G.PR., G.RG.; Writing - review and editing: G.PR., G.RG., A.CP., A.O.RM., M.A.RG., O.RM., M.G.GG., D.L.T., B.S., E.H.PA., C.R.HC., J.FR.; Funding acquisition: J.FR., M.A.RG., D.L.T., B.S., E.H.PA.; Resources: G.PR., G.RG., A.CP., A.O.RM., A.OM., M.G.GG., O.RM., M.A.RG., D.L.T., B.S., E.H.PA.; Data Curation: G.PR., G.RG., A.CP., A.O.RM., A.OM., M.G.GG., O.RM., M.A.RG., D.L.T.; Supervision: G.PR., G.RG., J.FR., C.R.HC. All authors reviewed the manuscript.

## Supplementary data

**Fig. SI1.**
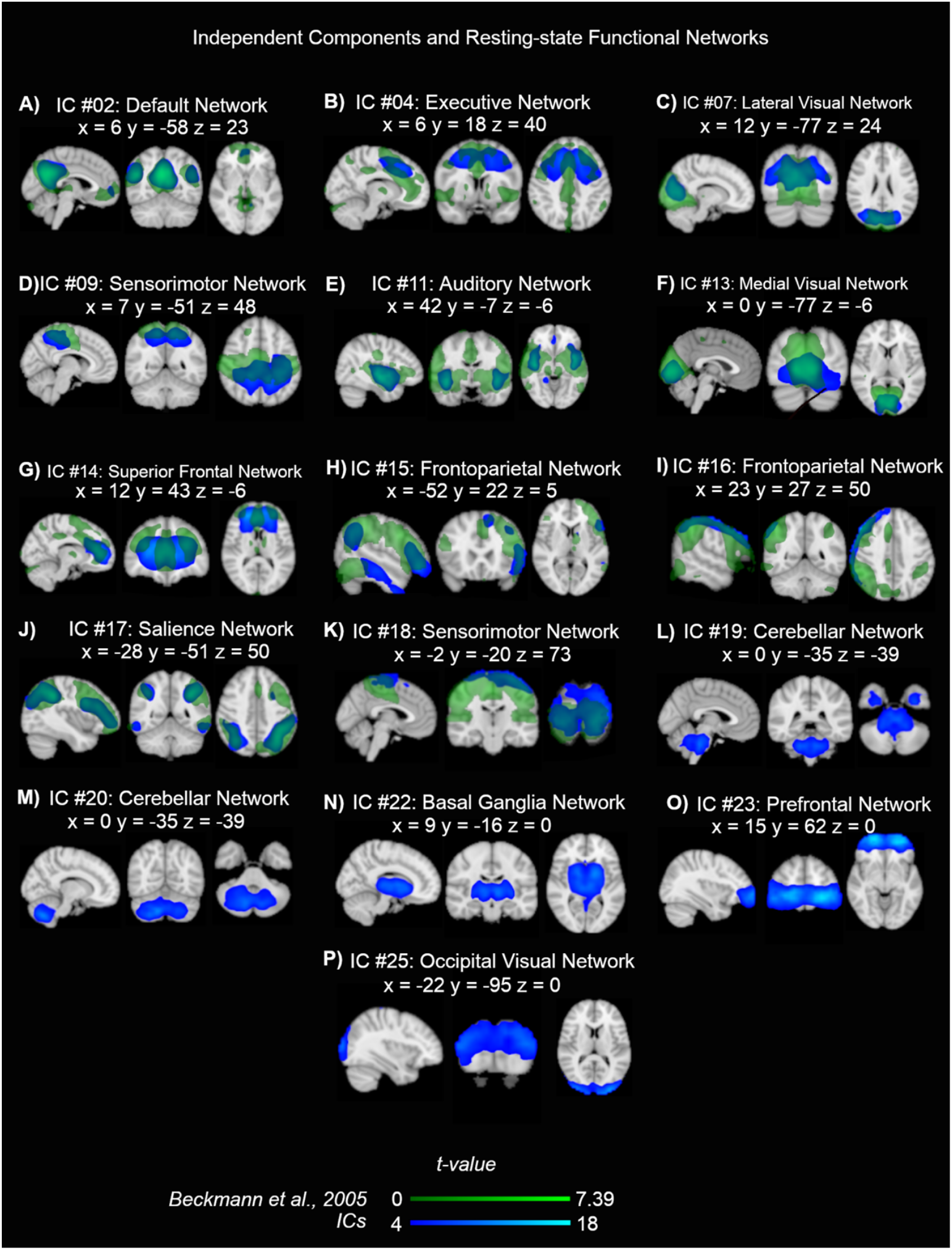
Sixteen Functional Networks were identified, after ICA, by overlapping a template of RSNs (blue and green color: A-K), and visual inspection (blue color: A-P)

## References

1 Matsuura T, Yamagata T, Burgess DL, Rasmussen A, Grewal RP, Watase K, et al. Large expansion of the ATTCT pentanucleotide repeat in spinocerebellar ataxia type 10. Nat Genet. 2000;26(2):191–4.

2 Teive HAG, Arruda WO. Cognitive dysfunction in spinocerebellar ataxias. Dement Neuropsychol. 2009;3(3):180–7.

3 Gatto EM, Gao R, White MC, Uribe Roca MC, Etcheverry JL, Persi G, et al. Ethnic origin and extrapyramidal signs in an Argentinean spinocerebellar ataxia type 10 family. Neurology. 2007;69(2):216–8.

4 Klockgether T, Mariotti C, Paulson HL. Spinocerebellar ataxia. Nat Rev Dis Prim. 2019;5(1):1–21.

5 Rasmussen A, Matsuura T, Ruano L, Yescas P, Ochoa A, Ashizawa T, et al. Clinical and genetic analysis of four Mexican families with spinocerebellar ataxia type 10. Ann Neurol. 2001;50(2):234–9.

6 Arruda WO, Meira AT, Ono SE, de Carvalho Neto A, Betting LEGG, Raskin S, et al. Volumetric MRI Changes in Spinocerebellar Ataxia (SCA3 and SCA10) Patients. Cerebellum. 2020;19(4):536–43.

7 Schmitz-Hübsch T, Du Montcel ST, Baliko L, Berciano J, Boesch S, Depondt C, et al. Scale for the assessment and rating of ataxia: Development of a new clinical scale. Neurology. 2006;66(11):1717–20.

8 Aguilar-Navarro SG, Mimenza-Alvarado AJ, Palacios-García AA, Samudio-Cruz A, Gutiérrez-Gutiérrez LA, Ávila-Funes JA. Validity and Reliability of the Spanish Version of the Montreal Cognitive Assessment (MoCA) for the Detection of Cognitive Impairment in Mexico. Rev Colomb Psiquiatr. 2018;47(4):237–43.

9 Larner AJ. Cognitive screening instruments: A practical approach. 2016. DOI: 10.1007/978-3-319-44775-9

10 Werner CJ, Dogan I, Saß C, Mirzazade S, Schiefer J, Shah NJ, et al. Altered resting-state connectivity in Huntington’s Disease. Hum Brain Mapp. 2014;35(6):2582–93.

11 Poudel GR, Egan GF, Churchyard A, Chua P, Stout JC, Georgiou-Karistianis N. Abnormal synchrony of resting state networks in premanifest and symptomatic Huntington disease: The IMAGE-HD study. J Psychiatry Neurosci. 2014;39(2):87– 96.

12 Pruim RHR, Mennes M, van Rooij D, Llera A, Buitelaar JK, Beckmann CF. ICA-AROMA: A robust ICA-based strategy for removing motion artifacts from fMRI data. Neuroimage. 2015;112:267–77.

13 Kairov U, Cantini L, Greco A, Molkenov A, Czerwinska U, Barillot E, et al. Determining the optimal number of independent components for reproducible transcriptomic data analysis. BMC Genomics. 2017;18(1):1–13.

14 Beckmann CF, DeLuca M, Devlin JT, Smith SM. Investigations into resting-state connectivity using independent component analysis. Philos Trans R Soc B Biol Sci. 2005;360(1457):1001–13.

15 Smith SM, Fox PT, Miller KL, Glahn DC, Fox PM, Mackay CE, et al. Correspondence of the brain’s functional architecture during activation and rest. Proc Natl Acad Sci U S A. 2009;106(31):13040–5.

16 Schimmelpfennig J, Topczewski J, Zajkowski W, Jankowiak-Siuda K. The role of the salience network in cognitive and affective deficits. Front Hum Neurosci. 2023;17(March):1–9.

17 Lee MH, Smyser CD, Shimony JS. Resting-state fMRI: A review of methods and clinical applications. Am J Neuroradiol. 2013;34(10):1866–72.

18 Raichle ME. Two views of brain function. Trends Cogn Sci. 2010;14(4):180–90.

19 Robinson S, Basso G, Soldati N, Sailer U, Jovicich J, Bruzzone L, et al. A resting state network in the motor control circuit of the basal ganglia. BMC Neurosci. 2009;10(Dm):1–14.

20 Catalino, M. P., Yao, S., Green, D. L., Laws, E. R., Golby, A. J., & Tie Y. Mapping cognitive and emotional networks in neurosurgical patients using resting-state functional magnetic resonance imaging. Neurosurg Focus. 2020;48(2). DOI: 10.1159/000444169.Carotid

21 Beckmann C, Mackay C, Filippini N, Smith S. Group comparison of resting-state FMRI data using multi-subject ICA and dual regression. Neuroimage. 2009;47:S148.

22 Nickerson LD, Smith SM, Öngür D, Beckmann CF. Using dual regression to investigate network shape and amplitude in functional connectivity analyses. Front Neurosci. 2017;11(MAR):1–18.

23 Diedrichsen J, Balsters JH, Flavell J, Cussans E, Ramnani N. A probabilistic MR atlas of the human cerebellum. Neuroimage. 2009;46(1):39–46.

24 Chirino-Pérez A, Vaca-Palomares I, Torres DL, Hernandez-Castillo CR, Diaz R, Ramirez-Garcia G, et al. Cognitive Impairments in Spinocerebellar Ataxia Type 10 and Their Relation to Cortical Thickness. Mov Disord. 2021;36(12):2910–21.

25 Hernandez-Castillo CR, Diaz R, Vaca-Palomares I, Torres DL, Chirino A, Campos-Romo A, et al. Extensive cerebellar and thalamic degeneration in spinocerebellar ataxia type 10. Park Relat Disord. 2019;66(May):182–8.

26 Cavanna AE, Trimble MR. The precuneus: A review of its functional anatomy and behavioural correlates. Brain. 2006;129(3):564–83.

27 Guo J, Jiang Z, Liu X, Li H, Biswal BB, Zhou B, et al. Cerebello-cerebral resting-state functional connectivity in spinocerebellar ataxia type 3. Hum Brain Mapp. 2023;44(3):927–36.

28 Rudolph S, Badura A, Lutzu S, Pathak SS, Thieme A, Verpeut JL, et al. Cognitive-Affective Functions of the Cerebellum. 2023;43(45):7554–64.

29 Siciliano L, Olivito G, Urbini N, Silveri MC, Leggio M. The rising role of cognitive reserve and associated compensatory brain networks in spinocerebellar ataxia type 2. J Neurol. 2023;270(10):5071–84.

30 Hoche F, Guell X, Vangel MG, Sherman JC, Schmahmann JD. The cerebellar cognitive affective/Schmahmann syndrome scale. Brain. 2018;141(1):248–70.

